# *In vivo* bioreactor matures iPSC-CMs for MLP disease modeling

**DOI:** 10.1101/2025.08.11.669708

**Authors:** Sean A. Murphy, Gunsik Cho, Pattraranee Limphong, Dong I. Lee, Chulan Kwon

## Abstract

Muscle LIM protein (MLP) is a critical regulator of cardiomyocytes (CMs) cytoarchitecture, and its deficiency results in late-onset dilated cardiomyopathy (DCM) in both mice and human. However, recapitulating this phenotype *in vitro* using induced pluripotent stem cell-derived cardiomyocytes (iPSC-CMs) has been challenging largely due to their immature state. Here, we generated MLP knockout (MLP-KO) mouse iPSCs and differentiated them into CMs. We then employed both conventional *in vitro* and *in vivo* transplantation approach using immunocompromised rat hearts to promote cardiomyocytes maturation. Our results showed that while *in vitro*-matured MLP-KO iPSC-CMs failed to exhibit disease phenotypes, *in vivo*-matured MLP-KO iPSC-CMs successfully recapitulated the hallmarks of DCM, including disrupted sarcomeric architecture and upregulation of atrial natriuretic peptide (ANP), closely mirroring disease progression observed in MLP-deficient mice. These findings demonstrate that the *in vivo* maturation environment is essential for the maturation of iPSC-derived cardiomyocytes to better model genetic cardiac diseases like DCM and provide valuable insights for future therapeutic strategies.

## Introduction

Mutation in the muscle LIM protein (MLP, also known as CSRP3) gene have been identified as critical contributors to both familial and sporadic forms of dilated cardiomyopathy (DCM), which is a major cause of heart failure and sudden cardiac death worldwide characterized by ventricular chamber enlargement and systolic dysfunction (Towbin et al., 2006). The etiology of DCM is multifactorial and heterogeneous, involving genetic mutations, viral infections, and idiopathic factors, thereby complicating both diagnosis and treatment strategies (McNally et al., 2013).

MLP is a striated muscle-specific protein that localizes to the Z-disc, where it plays a pivotal role in maintaining the structural integrity of the sarcomere and functions as a mechanosensor within cardiomyocytes (Arber et al., 1997; Knöll et al., 2002; Vafiadaki et al., 2015). Although MLP-deficient mice initially display normal cardiac morphology during early postnatal stages, they progressively develop phenotypic features characteristic of DCM, including ventricular dilation, systolic dysfunction, and heart failure, which closely recapitulate the progression seen in human patients with the condition (Knöll et al., 2002). Notably, at the molecular level, the absence of MLP leads to the disorganization of the sarcomeric Z-disc and the upregulation of fetal gene markers, such as atrial natriuretic peptide (ANP) and brain natriuretic peptide (BNP), which are molecular hallmarks of pathological cardiac remodeling (Heineke and Molkentin, 2006).

Induced pluripotent stem cells (iPSCs) have emerged as a revolutionary tool for modeling genetic cardiac diseases, allowing the derivation of patient- or mutation-specific cardiomyocytes *in vitro* (Moretti et al., 2010; Takahashi and Yamanaka, 2006). However, a significant challenge remains in the structural, functional, and transcriptomic maturation of iPSC-derived cardiomyocytes (iPSC-CMs), which often retain immature, fetal-like characteristics that limit their ability to replicate the structural and functional properties of adult cardiac disease phenotypes (Denning et al., 2016; Kannan and Kwon, 2020; Kannan et al., 2023; Murphy et al., 2021b, 2021a; Uosaki et al., 2015; Yang et al., 2014). Various strategies, including prolonged culture, electrical stimulation, and advance tissue engineering techniques, have been explored to enhance the maturation of iPSC-CMs and improve their adult-like characteristics (Lui et al., 2021; Nunes et al., 2013; Ronaldson-Bouchard et al., 2018; Shadrin et al., 2017). Nevertheless, these approaches have shown only limited success in achieving the desired adult-like properties (Kannan et al., 2021).

Recent studies have increasingly highlighted the essential role of *in vivo* environments in providing critical physiological stimuli for cardiomyocyte maturation, including factors that are often challenging to replicate effectively *in vitro*, such as mechanical load, neurohormonal signaling, and metabolic substrate availability (Funakoshi et al., 2016; Kadota et al., 2017). The transplantation of iPSC-CMs into animal models has been shown to enhance their structural and electrophysiological maturity, suggesting that *in vivo* maturation is crucial for accurately modeling late-onset cardiac diseases and capturing their complex pathophysiology (Cho et al., 2017a).

In this study, we aimed to generate MLP-KO mouse iPSCs and differentiate them into cardiomyocytes to establish a relevant model of genetic DCM. By comparing disease phenotypes under both *in vitro* and *in vivo* maturation conditions, we sought to determine if the physiological cues provided by the *in vivo* environment are necessary to fully recapitulate the pathological characteristics of DCM. This model not only offers insights into the mechanisms underlying MLP-related DCM but also establishes a foundation for future therapeutic screening, providing a platform for advancing treatment strategies in genetically characterized cardiac diseases. The understanding acquired from this research could inform strategies aimed at the development of advanced interventions for patients with genetic predispositions to DCM, ultimately leading to improved prognoses in this serious cardiovascular disease.

## Materials and Methods

### Animal work and Generation of MLP-KO iPSCs

Animal protocols were approved by the animal and care use committee at Johns Hopkins Medical Institutions. MLP-KO mice were obtained from the Jackson Laboratory. Tail-tip fibroblasts were isolated MLP-KO mice and cultured. Cells were reprogrammed using retroviral vectors encoding Klf4, Sox2, Oct4, and c-Myc. iPSC colonies were identified by morphology and alkaline phosphatase staining, and pluripotency was confirmed by immunofluorescence for Nanog and Oct4, as well as trilineage differentiation.

### Cardiac Progenitor Cell and Cardiomyocyte Differentiation and Maturation

The differentiation of iPSCs into CMs was performed as previously described (Takahashi and Yamanaka, 2006; Uosaki et al., 2012). Briefly, mouse iPSC were maintained in DMEM was supplemented with 15% fetal bovine serum, 0.1 mM 2-mercaptoethanol (Sigma-Aldrich), 0.1 mM sodium pyruvate (Invitrogen 0.1 mM non-essential amino acids, 2 mM L-glutamine (Invitrogen GlutaMAX), and 100 Units/ml leukemia inhibitory factor (LIF, Millipore), 1 µM PD0325901, and 3 µM CHIR99021. For cardiac progenitor cell differentiation, iPSCs were dissociated to single cells and transferring 75,000 per mL to ultra-low attachment plates (Corning), and cultured in IMDM/Ham-F12 (Cellgro) supplemented with various factors including 2 mM L-glutamine (Invitrogen GlutaMAX), N2, 0.05% bovine serum albumin, B27, 5 ng/ml L-ascorbic acid (Sigma-Aldrich), and α-monothioglycerol (Sigma-Aldrich), 1X penicillin/streptomycin (Invitrogen). On day 7 after differentiation, CPCs were sorted. Differentiated CMs were matured either *in vitro* (2D culture for 30 days) or *in vivo* by transplantation into immunocompromised rat hearts, with CMs labeled with GFP for lineage tracking (Cho et al., 2017b).

### Cell Sorting and Transplantation

On day 7 of differentiation, cells were dissociated and resuspended in PBS with 20 mM HEPES, 0.1 % FBS, and 1mM EDTA. Fluorescence-activated cell sorting was performed with Sony SH800 to isolate GFP. Nude rats (Charles River Laboratories) were intracardially injected with 2×10^5^ CPCs IMDM with 1:60 Matrigel through a glass micropipette held by a micromanipulator. Prior to injection, neonatal rats were anesthetized on ice and had a hole surgically opened between the 4^th^ and 5^th^ rib. 3M Vetbond tissue adhesion was applied to the hole before letting the rat pups recover on a heating pad.

### Immunofluorescence, Histological Analysis, and Imaging

To confirm tri lineage potential, embryoid bodies were formed and stained for germ layer markers: α-SMA (mesoderm), AFP (endoderm), and GFAP (ectoderm). Heart and graft tissues were fixed with 4% paraformaldehyde and in optimal cutting temperature (OCT) compound before cryosectioning. They were stained with antibodies against ANP, α-actinin (Sigma A7732), and wheat germ agglutinin conjugated to Alexa Fluor 568. Nuclei were counterstained with DAPI. Fluorescence imaging was performed using confocal microscopy on a Leica DM2500 microscope with a 40x oil-immersion lens. ANP expression intensity was quantified using ImageJ software.

### qPCR

The Trizol protocol was used for RNA Isolation. RNA was reverse transcribed using high-capacity cDNA kit (Applied Biosystems). qPCR amplification used Syber Select qPCR master mix (Thermo Fisher) on a CFX96 (Biorad).

### Statistical Analysis

Statistical analysis was performed using paired Student’s t-test; p-values <0.05 were considered significant.

## Results

### Generation of MLP Knockout iPSCs Line

To investigate if the DCM phenotype could be replicated in an iPSC model, we generated MLP-deficient miPSCs from MLP knockout (KO) mice via expression of the reprogramming factors Klf4, Sox2, Oct4, and c-Myc. The pluripotency of the resulting iPSCs was confirmed through immunostaining for the pluripotency markers NANOG and SSEA4 (**Figure 1A**), indicating successful reprogramming. Further, to assess their developmental potential, these MLP-KO miPSCs were subjected to spontaneous differentiation. The expression of lineage markers such as α-Fetoprotein (endoderm), α-Smooth Muscle Actin (mesoderm), and Glial Fibrillary Acidic Protein (ectoderm) demonstrated their ability to differentiate into all three germ layers, confirming the multipotency of iPSCs (**Figure 1B**).

**Figure 1:**
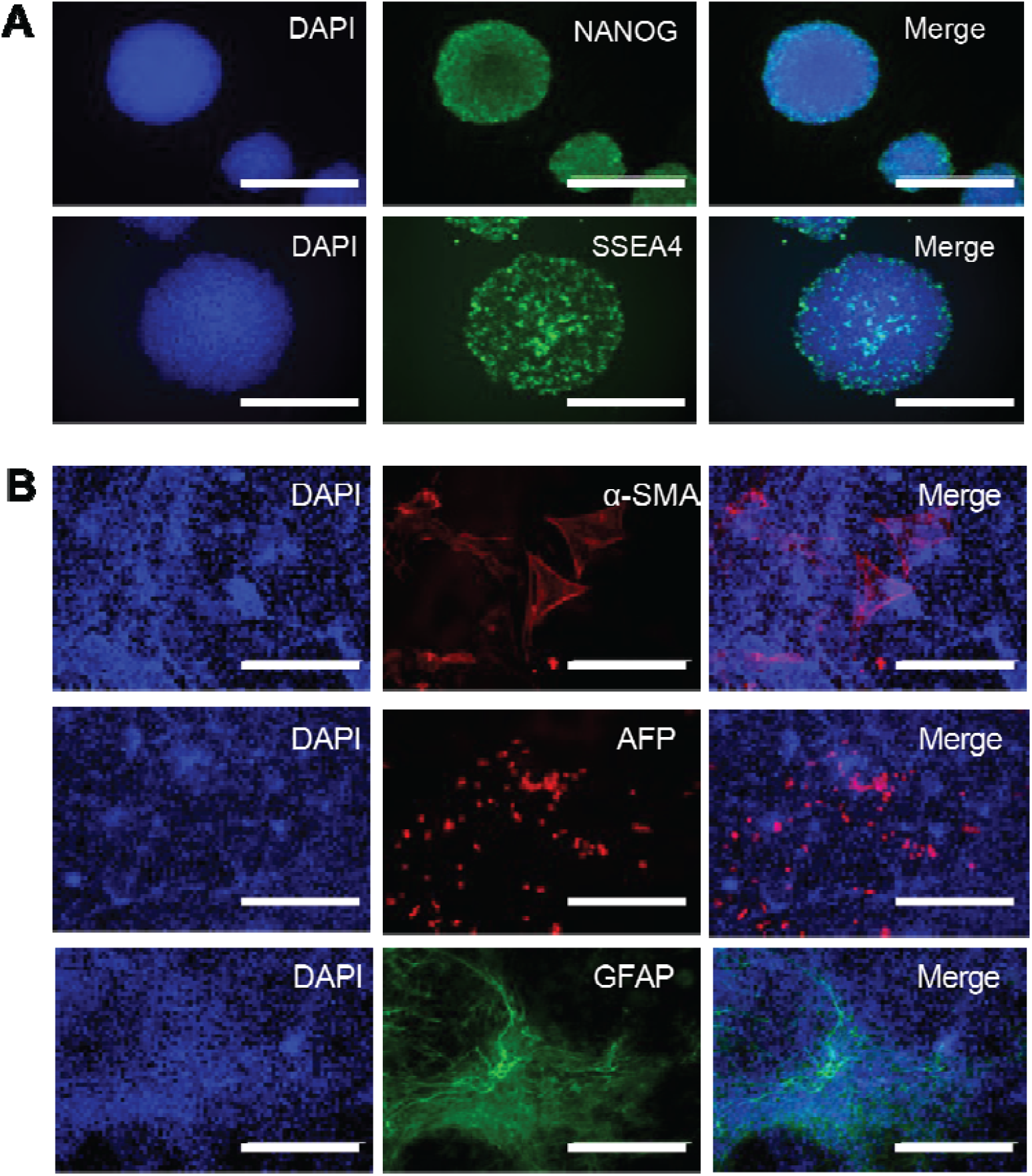
Generation of iPSCs from MLP KO Mice. **(A)** MLP KO mouse derived iPSC colonies immunostained for pluripotency markers NANOG and SSEA4. **(B)** MLP KO iPSC colonies stained with pluripotent stem cell markers. (d) Differentiating MLP-iPSCs stained with three germ layers markers alpha smooth muscle actin (α-SMA; mesoderm), alpha-fetoprotein (AFP; endoderm) and Glial Fibrillary Acidic Protein (GFAP; ectoderm).

### MLP Knockout Mouse Displays Dilated Cardiomyopathy

Consistent with previous report (Knöll et al., 2002), mice lacking MLP developed the characteristic manifestations of DCM, a late-onset cardiac disease (**Figure 2A)**. Although MLP-deficient mice exhibited normal morphology during the early postnatal period, they subsequently developed an abnormal cardiac phenotype, including depressed function, disorganized myocyte sarcomeres, and upregulation of the hypertrophy marker atrial natriuretic peptide (ANP) (**Figure 2B**).

**Figure 2.**
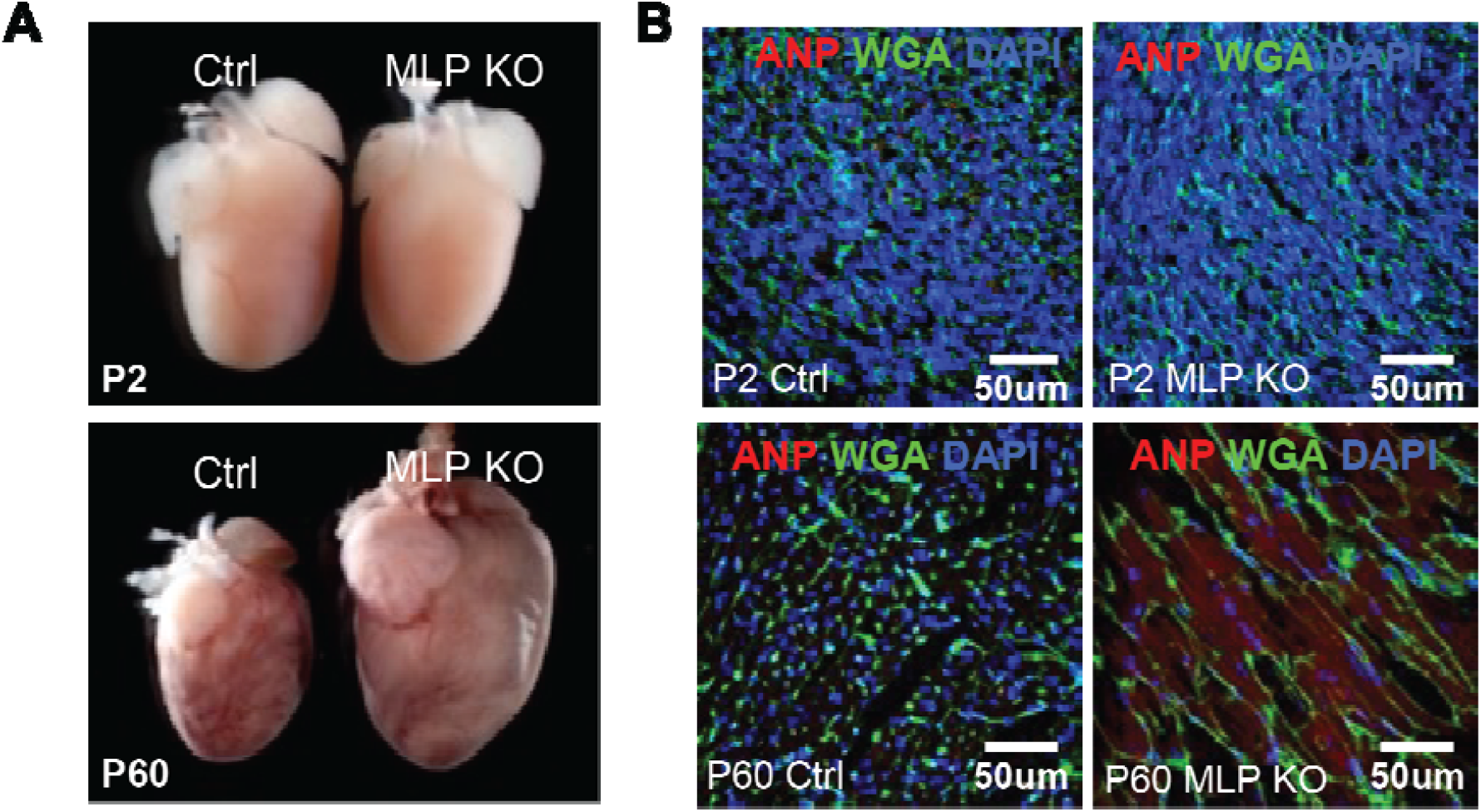
MLP KO heart morphology and ANP expression. **(A)** Whole heart comparisons of control and MLP KO mice at indicated stages. n=3 mice. **(B)** ANP (red) and WGA (green) staining of control and MLP KO heart sections at indicated stages.

### In vivo Maturation System Allows Modeling Dilated Cardiomyopathy

To assess the functional maturation of MLP-KO iPSCs, we injected them into the hearts of neonatal immunodeficient rat. Before injection, nuclei were labeled with GFP and iPSCs were differentiated into CMs. As a control, MLP-KO miPSC-CMs that were cultured *in vitro* were evaluated. The *in vitro*-cultured MLP-KO miPSC-CMs remained morphologically indistinguishable from control miPSC-CMs, and there was no difference in ANP levels (**Figure 3A**). However, the *in vivo*-matured MLP-KO miPSC-CMs showed a diffused α-actinin expression, closely recapitulating the *in vivo* phenotype observed in DCM (**Figure 3B**). This was accompanied by high levels of ANP expression in P60 *in vivo*-matured miPSC-CMs, while P2 miPSC-CMs showed no significant difference compared to control miPSC-CMs (**Figure 4A**). Furthermorer, *in vivo*-matured MLP KO miPSC-CMs displayed a significant loss of sarcomeric ultrastructure, which is consistent with the DCM phenotype (**Figure 4B**). These findings suggest that the *in vivo* PSC maturation system can be used for modeling late-onset heart muscle disease and provide critical insights into DCM progression and mechanisms.

**Figure 3.**
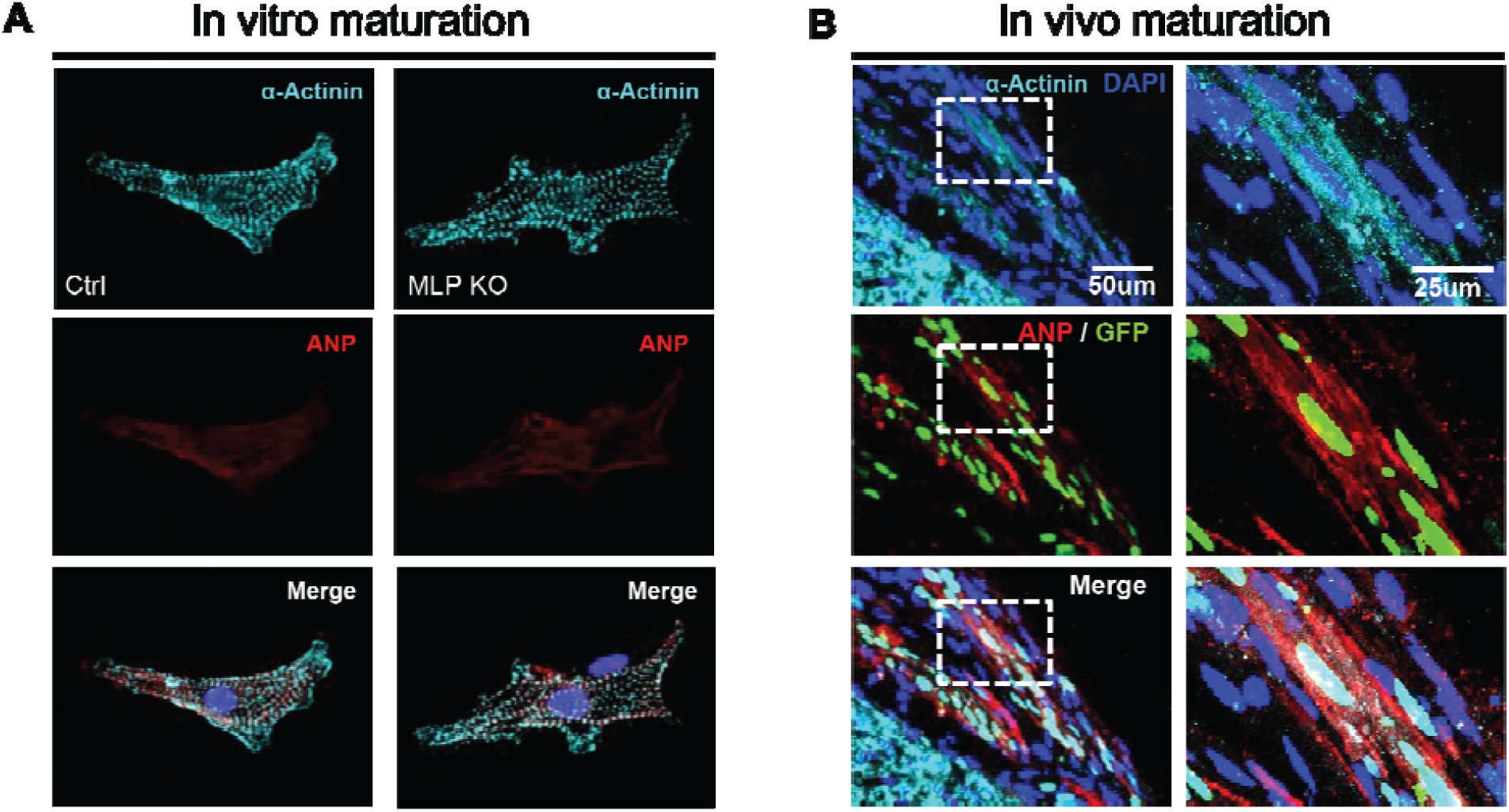
*In vivo* maturation of MLP-deficient miPSC-CMs. **(A)** *In vitro*-matured control and MLP-KO iPSC-CMs stained with ANP (red) and α-Actinin (cyan). **(B)** *In vivo*-matured MLP-KO iPSC-CMs (labeled with nuclear GFP) stained with α-Actinin and ANP. n=5 Rats. Right panel shows enlargement of boxed areas (left). DAPI (blue) was used to counterstain nuclei. Ctrl, control; M, month; p, postnatal day.

**Figure 4.**
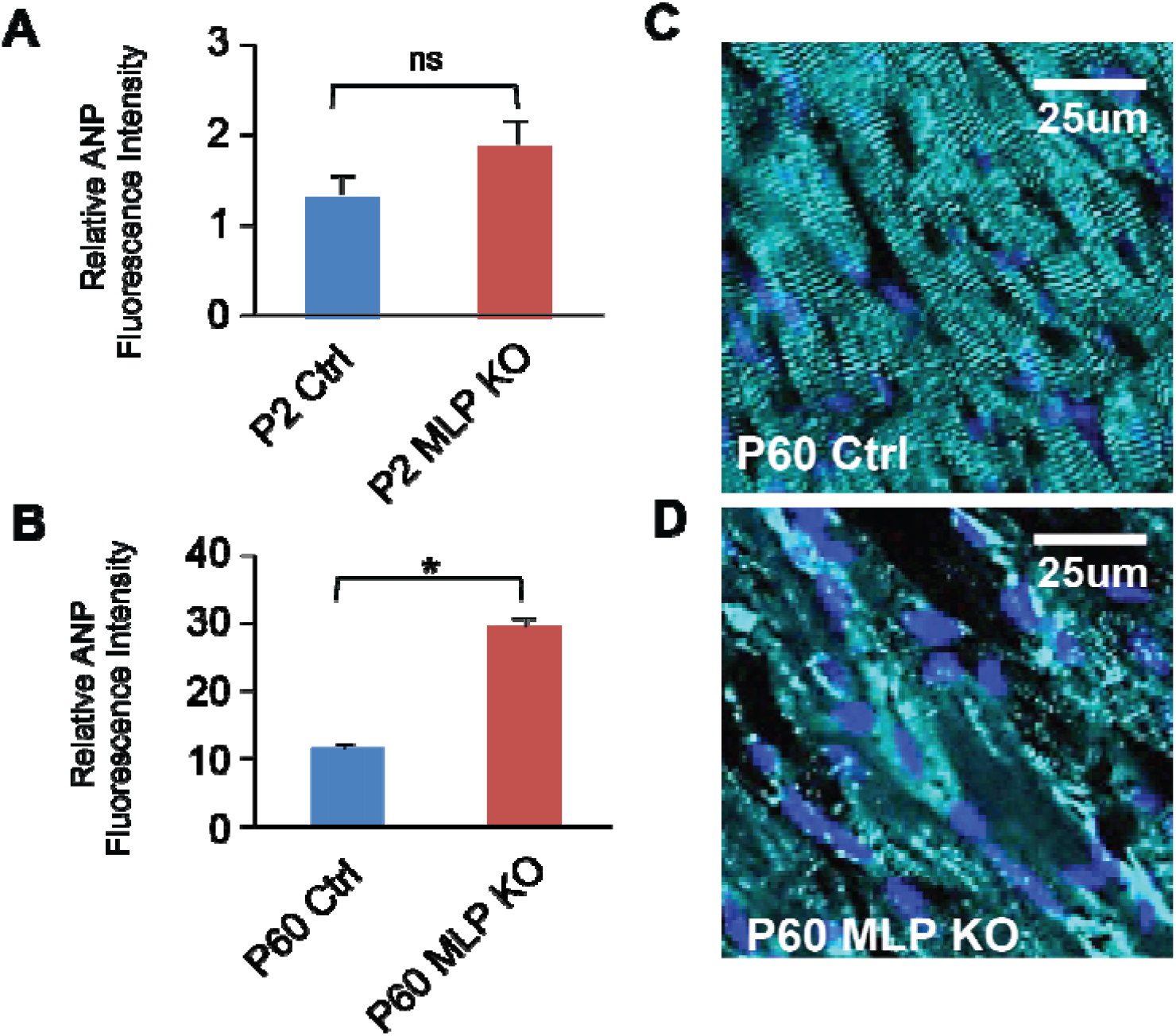
Increased ANP and disorganized sarcomeres in MLP-deficient iPSC-CMs after *in vivo* Maturation. **(A)** Relative ANP expression intensity in control and mutant hearts at P2 and (**B**) P60, measured by image J. Data are mean ± SD; *p<0.05. *ns*, not significant (p>0.05). p values were determined using the paired Student *t* test. **(C)** α-Actinin (cyan) staining in control and (**D**) MLP-KO CMs hearts showing disorganized sarcomeric structure.

## Discussion

In this study, we provide compelling evidence that *in vivo* maturation is crucial for modeling late-onset generic cardiac diseases, such as DCM, using iPSC-derived cardiomyocytes. Consistent with previous reports, MLP-KO mice did not show structural or functional cardiac abnormalities during early models that recapitulate human DCM phenotypes, including ventricular dilation and systolic dysfunction, as they mature (Knöll et al., 2002). Despite this, our *in vitro*-differentiated MLP-KO iPSC-CMs failed to exhibit key disease characteristics such as sarcomeric disarray, impaired contractility, or upregulation of fetal genes, likely due to their immature phenotype and the absence of critical biomechanical and neurohormonal stimuli present *in vivo* (Denning et al., 2016; Yang et al., 2014). This is consistent with previous reports highlighting the limitations of current *in vitro* maturation protocols, which often cannot produce functionally mature, adult-like cardiomyocytes capable of mimicking late-onset cardiac disease phenotypes (Nunes et al., 2013; Ronaldson-Bouchard et al., 2018).

In contrast, our study demonstrated that MLP-KO iPSC-CMs matured *in vivo* showed sarcomeric disorganization and upregulated ANP expression, both of which are hallmarks of pathological remodeling found in MLP-deficient mice and human DCM (Heineke and Molkentin, 2006; Vafiadaki et al., 2015). These findings highlight the inadequacy of current *in vitro* maturation protocols and stress the importance of *in vivo* physiological cues, such as mechanical stress, neurohormonal signaling, and optimal metabolic substrate availability, for the manifestation of disease phenotypes in genetically altered cardiomyocytes (Funakoshi et al., 2016; Kadota et al., 2017; Ronaldson-Bouchard et al., 2018). Our results indicate that for diseases like DCM, where clinical symptoms and cellular pathology emerge progressively under physiological stress, *in vitro* models may lack sufficient maturation stimuli to fully express the disease phenotypes.

This work supports the notion that *in vivo* maturation strategies should be an integral component of iPSC-based disease modeling, particularly for disorders characterized by delayed onset or complex structural components. The successful application of an *in vivo* maturation platform, as demonstrated in our study, holds great promise for future applications across various areas including disease modeling, drug screening, and regenerative therapy. By incorporating *in vivo* maturation techniques, it may be possible to unveil therapeutic targets that might remain undetected in immature *in vitro* systems and assess pharmacological interventions in a context that more accurately reflects the disease state.

Furthermore, our finding emphasizes the relevance of employing a comprehensive approach to investigating the cellular and molecular mechanisms that underlie genetic cardiac diseases such as DCM. This is crucial not only for advancing fundamental scientific knowledge but also for developing targeted therapeutic that improve patients outcomes. Ultimately, by establishing a reliable model that incorporates the complexities of *in vivo* physiology, we aim to generate more effective treatment strategies for patients with genetic predispositions to DCM, contributing to the broader landscape of cardiovascular medicine.

## Acknowledgement

This work was supported by grants from NHLBI/NIH (R01HL156947, R01HL171205) and AHA (23TPA1058685).

## Author Contributions

G.C., S.M., D.I.L. designed and carried out this work. P.L. performed iPSCs differentiation. D.I.L., C.K. designed and supervised this work and wrote the manuscript with S.M.

## Figures

**Figure S1.**
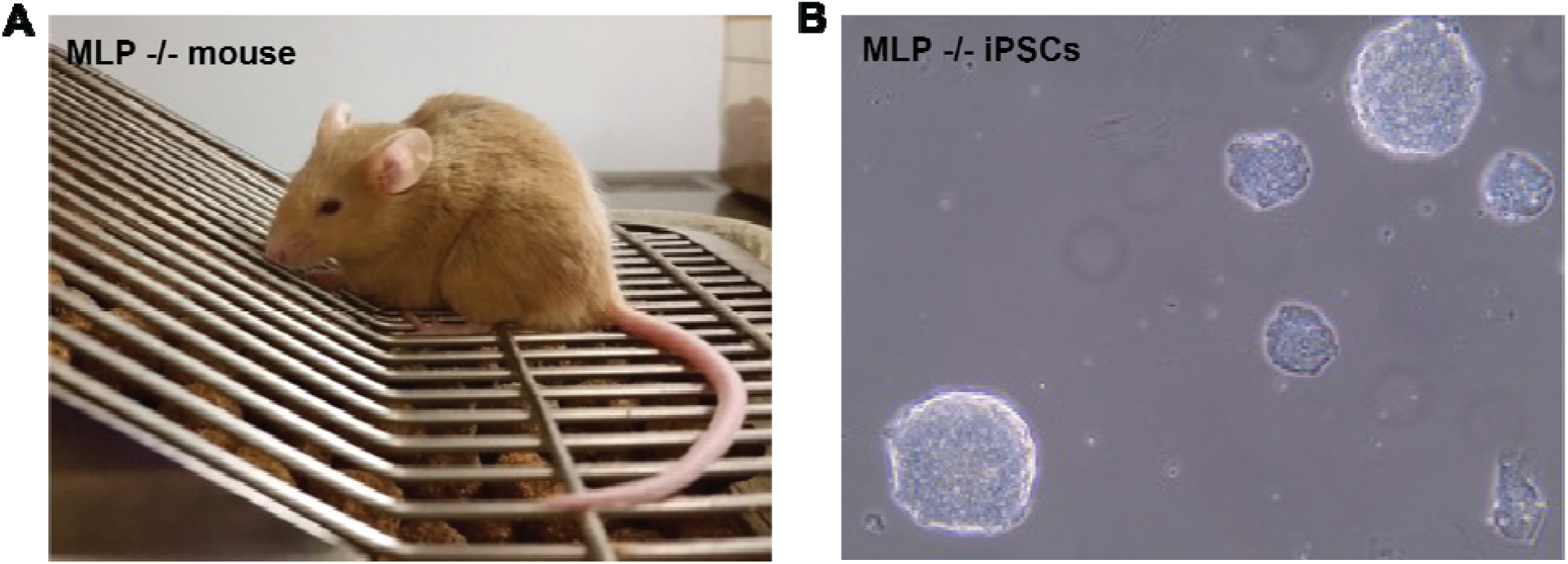
MLP knockout iPSC line. **(A)** MLP knockout mouse. **(B)** iPSCs form round colonies.

## Reference

Arber, S., Hunter, J.J., Ross, J., Hongo, M., Sansig, G., Borg, J., Perriard, J.C., Chien, K.R., and Caroni, P. (1997). MLP-deficient mice exhibit a disruption of cardiac cytoarchitectural organization, dilated cardiomyopathy, and heart failure. Cell 88, 393–403. 10.1016/s0092-8674(00)81878-4.

Cho, G.-S., Lee, D.I., Tampakakis, E., Murphy, S., Andersen, P., Uosaki, H., Chelko, S., Chakir, K., Hong, I., Seo, K., et al. (2017a). Neonatal Transplantation Confers Maturation of PSC-Derived Cardiomyocytes Conducive to Modeling Cardiomyopathy. Cell Reports 18, 571–582. 10.1016/j.celrep.2016.12.040.

Cho, G.-S., Tampakakis, E., Andersen, P., and Kwon, C. (2017b). Use of a neonatal rat system as a bioincubator to generate adult-like mature cardiomyocytes from human and mouse pluripotent stem cells. Nat Protoc 12, 2097–2109. 10.1038/nprot.2017.089.

Denning, C., Borgdorff, V., Crutchley, J., Firth, K.S.A., George, V., Kalra, S., Kondrashov, A., Hoang, M.D., Mosqueira, D., Patel, A., et al. (2016). Cardiomyocytes from human pluripotent stem cells: From laboratory curiosity to industrial biomedical platform. Biochim Biophys Acta 1863, 1728–1748. 10.1016/j.bbamcr.2015.10.014.

Funakoshi, S., Miki, K., Takaki, T., Okubo, C., Hatani, T., Chonabayashi, K., Nishikawa, M., Takei, I., Oishi, A., Narita, M., et al. (2016). Enhanced engraftment, proliferation, and therapeutic potential in heart using optimized human iPSC-derived cardiomyocytes. Sci Rep 6, 19111. 10.1038/srep19111.

Heineke, J., and Molkentin, J.D. (2006). Regulation of cardiac hypertrophy by intracellular signalling pathways. Nat Rev Mol Cell Biol 7, 589–600. 10.1038/nrm1983.

Kadota, S., Pabon, L., Reinecke, H., and Murry, C.E. (2017). In Vivo Maturation of Human Induced Pluripotent Stem Cell-Derived Cardiomyocytes in Neonatal and Adult Rat Hearts. Stem Cell Reports 8, 278–289. 10.1016/j.stemcr.2016.10.009.

Kannan, S., and Kwon, C. (2020). Regulation of cardiomyocyte maturation during critical perinatal window. J Physiol 598, 2941–2956. 10.1113/JP276754.

Kannan, S., Farid, M., Lin, B.L., Miyamoto, M., and Kwon, C. (2021). Transcriptomic entropy benchmarks stem cell-derived cardiomyocyte maturation against endogenous tissue at single cell level. PLoS Comput Biol 17, e1009305. 10.1371/journal.pcbi.1009305.

Kannan, S., Miyamoto, M., Zhu, R., Lynott, M., Guo, J., Chen, E.Z., Colas, A.R., Lin, B.L., and Kwon, C. (2023). Trajectory reconstruction identifies dysregulation of perinatal maturation programs in pluripotent stem cell-derived cardiomyocytes. Cell Rep 42, 112330. 10.1016/j.celrep.2023.112330.

Knöll, R., Hoshijima, M., Hoffman, H.M., Person, V., Lorenzen-Schmidt, I., Bang, M.-L., Hayashi, T., Shiga, N., Yasukawa, H., Schaper, W., et al. (2002). The cardiac mechanical stretch sensor machinery involves a Z disc complex that is defective in a subset of human dilated cardiomyopathy. Cell 111, 943–955. 10.1016/s0092-8674(02)01226-6.

Lui, C., Chin, A.F., Park, S., Yeung, E., Kwon, C., Tomaselli, G., Chen, Y., and Hibino, N. (2021). Mechanical stimulation enhances development of scaffold-free, 3D-printed, engineered heart tissue grafts. J Tissue Eng Regen Med 15, 503–512. 10.1002/term.3188.

McNally, E.M., Golbus, J.R., and Puckelwartz, M.J. (2013). Genetic mutations and mechanisms in dilated cardiomyopathy. J Clin Invest 123, 19–26. 10.1172/JCI62862.

Moretti, A., Bellin, M., Welling, A., Jung, C.B., Lam, J.T., Bott-Flügel, L., Dorn, T., Goedel, A., Höhnke, C., Hofmann, F., et al. (2010). Patient-specific induced pluripotent stem-cell models for long-QT syndrome. N Engl J Med 363, 1397–1409. 10.1056/NEJMoa0908679.

Murphy, S.A., Miyamoto, M., Kervadec, A., Kannan, S., Tampakakis, E., Kambhampati, S., Lin, B.L., Paek, S., Andersen, P., Lee, D.-I., et al. (2021a). PGC1/PPAR drive cardiomyocyte maturation at single cell level via YAP1 and SF3B2. Nat Commun 12, 1648. 10.1038/s41467-021-21957-z.

Murphy, S.A., Chen, E.Z., Tung, L., Boheler, K.R., and Kwon, C. (2021b). Maturing heart muscle cells: Mechanisms and transcriptomic insights. Semin Cell Dev Biol 119, 49–60. 10.1016/j.semcdb.2021.04.019.

Nunes, S.S., Miklas, J.W., Liu, J., Aschar-Sobbi, R., Xiao, Y., Zhang, B., Jiang, J., Massé, S., Gagliardi, M., Hsieh, A., et al. (2013). Biowire: a platform for maturation of human pluripotent stem cell-derived cardiomyocytes. Nat Methods 10, 781–787. 10.1038/nmeth.2524.

Ronaldson-Bouchard, K., Ma, S.P., Yeager, K., Chen, T., Song, L., Sirabella, D., Morikawa, K., Teles, D., Yazawa, M., and Vunjak-Novakovic, G. (2018). Advanced maturation of human cardiac tissue grown from pluripotent stem cells. Nature 556, 239–243. 10.1038/s41586-018-0016-3.

Shadrin, I.Y., Allen, B.W., Qian, Y., Jackman, C.P., Carlson, A.L., Juhas, M.E., and Bursac, N. (2017). Cardiopatch platform enables maturation and scale-up of human pluripotent stem cell-derived engineered heart tissues. Nat Commun 8, 1825. 10.1038/s41467-017-01946-x.

Takahashi, K., and Yamanaka, S. (2006). Induction of pluripotent stem cells from mouse embryonic and adult fibroblast cultures by defined factors. Cell 126, 663–676. 10.1016/j.cell.2006.07.024.

Towbin, J.A., Lowe, A.M., Colan, S.D., Sleeper, L.A., Orav, E.J., Clunie, S., Messere, J., Cox, G.F., Lurie, P.R., Hsu, D., et al. (2006). Incidence, Causes, and Outcomes of Dilated Cardiomyopathy in Children. JAMA 296, 1867–1876. 10.1001/jama.296.15.1867.

Uosaki, H., Andersen, P., Shenje, L.T., Fernandez, L., Christiansen, S.L., and Kwon, C. (2012). Direct contact with endoderm-like cells efficiently induces cardiac progenitors from mouse and human pluripotent stem cells. PLoS One 7, e46413. 10.1371/journal.pone.0046413.

Uosaki, H., Cahan, P., Lee, D.I., Wang, S., Miyamoto, M., Fernandez, L., Kass, D.A., and Kwon, C. (2015). Transcriptional Landscape of Cardiomyocyte Maturation. Cell Rep 13, 1705–1716. 10.1016/j.celrep.2015.10.032.

Vafiadaki, E., Arvanitis, D.A., and Sanoudou, D. (2015). Muscle LIM Protein: Master regulator of cardiac and skeletal muscle functions. Gene 566, 1–7. 10.1016/j.gene.2015.04.077.

Yang, X., Pabon, L., and Murry, C.E. (2014). Engineering adolescence: maturation of human pluripotent stem cell-derived cardiomyocytes. Circ Res 114, 511–523. 10.1161/CIRCRESAHA.114.300558.

